# Rapid quantitative assessment of small molecule leakage from microdroplets by flow cytometry and improvement of fluorophore retention in biochemical assays

**DOI:** 10.1101/2023.04.23.538007

**Authors:** Anastasia Zinchenko, Sean R. A. Devenish, Florian Hollfelder

## Abstract

Microdroplets are compartments made in the laboratory that allow the miniaturisation of chemical and biological experiments to the femto- to picolitre scale, replacing the classical test tube with a droplet. Ideally containment of the contents of individual droplets would be perfect, but in reality this situation rarely occurs. Instead the leaking of molecules even from intact droplets presents a challenge to the success of miniaturisation and must be assessed on a case-by-case basis. We now present a new method for quantitative determination of leakage: a sheath fluid-free flow cytometer (Guava EasyCyte) is used to directly determine the fluorescence of water-in-oil droplets as a function of time. We validate this method by demonstrating that this assessment of leakage provides a framework for experimental improvements that reduce the leakage of two widely used fluorophores. A 40-fold better retention compared to current protocols is achieved for resorufin with an optimized mix (oil: FC-70, surfactant: 0.1% w/w AZ900C, additive: 1% BSA) to maintain useful retention for up to 130 hours. Likewise leakage of the fluorophore methylumbelliferone is reduced by 75-fold. The availability of a method to quantitate leakage quickly for a variety of experimental conditions will facilitate future applications of droplet-based experiments (e.g. in directed evolution or diagnostics), aid miniaturisation of lab-scale assays into this format, and improve the degrees of freedom in setting up such ultrahigh-throughput experiments.

## Introduction

Future developments in biochemical assay technology will need to deliver both miniaturized sample volumes and increased rates of analysis to enable ever-larger screening campaigns with ultrahigh throughput. Both of these goals can be fulfilled by encapsulating (bio)chemical assays in water-in-oil microdroplets made in microfluidic devices^1–2^ or open systems.^3^ Conducting reactions in microdroplets is extremely reagent-saving, with assay volumes in the pico-to femtoliter range. Droplet-based high-throughput screening methods facilitate analysis of extraordinary sample sizes with throughputs in the tens of kHz range implying a capacity of up to over 100 million independent reactions in a working day.^1–2, 4–9^ This type of screening is advantageous in particular for diagnostic applications,^10–13^ or to identify efficient catalytic systems,^9, 14–15^ e.g. by interrogating protein libraries in directed evolution,^16–21^ and functional metagenomics,^22–24^ and in strain selections^25–26^ or bioprospecting.^27^ In each case positive hits in samples, libraries and populations are expected to be rare and must be reliably detected.^28^

In order to perform assays with a useful readout, a prime concern is to maintain the integrity of the droplet, avoiding fusion or other ‘droplet failure modes’.^29–30^ Secondly the droplets should not shrink or expand uncontrolledly.^31–32^ Finally small molecules, should not leave their droplet compartments: if they can, the assumption that each droplet is a unique microreactor is compromised or invalidated. Such leakage can happen after breakdown of the droplet integrity,^29^ but also – less obvious and thus harder to detect – between droplets that maintain their stability and look structurally intact by visual inspection. This last issue of small molecule transfer between intact droplets is addressed in this work (and referred to as ‘leakage’). It is well known^33–42^ that fluorescent reporter groups for processes studied in droplets are prone to transfer between them. This includes the fluorescent reaction products of many commercially available fluorogenic substrates for enzyme-catalysed reactions, as many of the most widely used small fluorescent molecules are hydrophobic, and similar problems may arise for conjugates detected by fluorescence anisotropy ^43^. Their low solubility in aqueous solution favors escape into the oil phase and transfer between droplets by diffusion. The use of fluorinated oils - designed to act as a ‘third phase’ with hydrophobic *and* lipophobic properties^44^ - instead of the mineral or silicon oils originally used in microdroplet research helps to reduce such leakage,^45–46^ but does not eliminate it completely. Exchange of fluorophore between droplets diminishes the signal difference between droplets containing biocatalysts with different activities. The lack of precise quantification of reaction product (or the product of a subsequent coupled assay reaction) thus hampers identification and selection of the most active variants in libraries of biocatalysts. Even though such leakage is a familiar practical shortcoming of microdroplet experiments and severely limits the range of bioassays that can be carried out, practical methods to enable empirical testing in the process of setting up novel assays are needed to obtain a quantitative measure of leakage will facilitate countering this problem in the future.

A recently introduced method elegantly employs electrospray ionization mass spectrometry (ESI) to investigate small molecule transfer between droplets.^47^ However, most previous studies of small molecule transfer from microdroplets have typically relied on either microscope imaging or fluorescence measurement on chip.^33–35, 41^ Recording microscope images is experimentally straightforward, but image analysis is time consuming. This means that the number of droplets that can be analysed in a reasonable timeframe is low (typically hundreds to thousands at best).^48^ Furthermore, the measurements obtained tend to be imprecise due to the imperfect lighting of typical microscope set-ups.^49^ On the other hand, analysis of microdroplet fluorescence on a microfluidic chip is technically challenging, requiring specialized equipment and knowledge in microfluidics, optics and electronics.^17, 50^ Such technology is readily available in specialist microengineering laboratories but not normally in biochemical research environments. We now introduce a method to rapidly measure small molecule transfer under a wide variety of conditions (e.g. with different oil phases, surfactant types, reagent concentrations or medium pH). The availability of this method will add to the surprisingly few systematic studies^29, 33, 51^ available and ultimately provide a basis for drawing up general guidelines for minimizing small molecule transfer based on quantitative data, as exemplified by a predictive tool based on principle component analysis.^47^ The availability of such guidelines for the choice of surfactant/oil combinations and feasibility of assays involving small molecules, would facilitate implementation of improvements on protocols for microdroplet experiments.

In the present work we address this shortcoming by developing an analytical method for accurate determination of the rate and extent of small molecule transfer from microdroplets, and then apply this method to a systematic exploration of the dependence of small molecule transfer on the fluorous oil carrier, the surfactant^30^ and the makeup of the compartmentalized aqueous phase (Fig. 1). By examining these key parameters we hope to derive broadly applicable strategies for preventing or minimizing small molecule transfer from microdroplets. The analytical method developed here exploits flow cytometry as a readily accessible, accurate and high throughput analytical technique. Conventional flow cytometer instruments built for analysis of biological cells utilize aqueous sheath fluid as a carrier phase and so do not allow analysis of water-in-oil emulsion droplets due to their oil carrier phase. Water-in-oil emulsion droplets can be transformed into the aqueous format of water-oil-water double emulsions before FACS analysis,^52–56^ but leakage is evident and often faster than from single emulsions.^52^ If emulsions of different content are emulsified in two separate steps and mixed as double emulsions, the fluorophore is more likely to leak into the aqueous carrier phase than into the neighbouring droplets, confounding collection of accurate data on exchange *via* the oil phase that is relevant in a typical biocatalyst screening experiment. Thus, we took advantage of the capacity of a microcapillary flow cytometry system (Guava EasyCyte 6HT-2L) to directly analyze water-in-oil droplets for size and fluorescence. In contrast to traditional flow cytometers this system does not use a sheath fluid and so oil can be used directly as a carrier phase. This facility enabled us to perform an accurate, simple and rapid measurement of small molecule transfer (by defining the parameter % leakage) at various time points under different conditions and thereby identify conditions that reduce the leakage of two important fluorophores, resorufin **1** and methylumbelliferone **2,** by almost two orders of magnitude.

**Figure 1.**
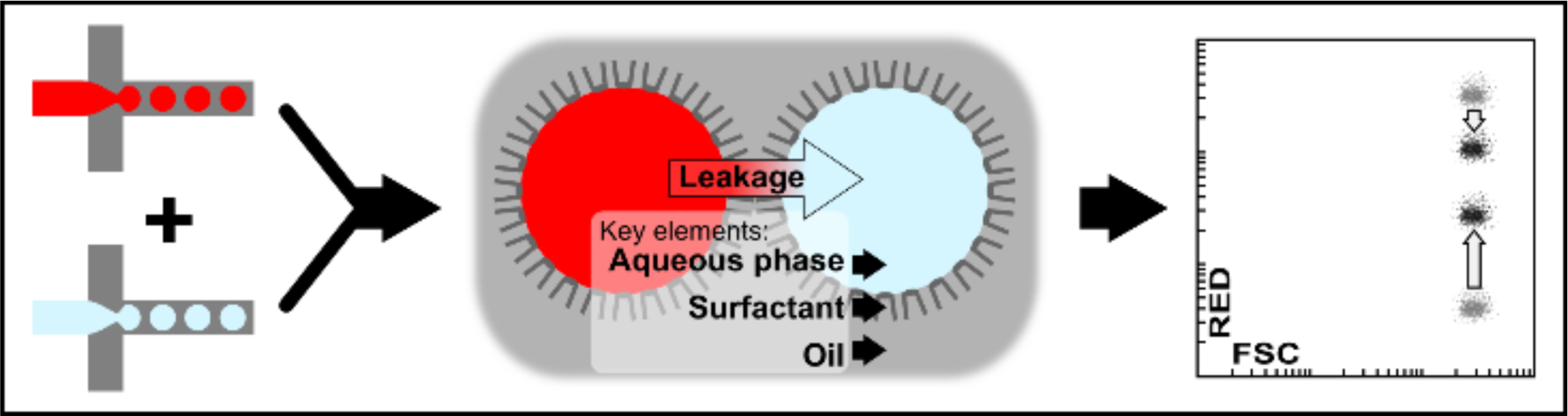
Schematic view of a methodology to assess small molecule leaking between droplets. Two populations of monodisperse droplets – one containing and one lacking a fluorophore - were produced in microfluidic flow focusing devices and mixed. During incubation fluorophore transfer between droplets can take place. The extent of such leakage was then assessed by sheath fluid-free flow cytometry for multiple time points. A typical outcome of such an analysis is illustrated by a leaking experiment with resorufin **1**: two initially distinct droplet populations assume more and more similar fluorescence values, suggesting exchange of the fluorophore (oil phase: HFE7500, surfactant: 1% AZ900C (w/w), time: superposed data for 0 and 4 hours). The primary determinants of the rate of leakage were found to be the oil, the surfactant concentration and the nature of aqueous phase additives. Such experiments provide means to optimize the experimental conditions for biological and chemical experiments in droplets with minimized small molecule leakage.

## Results and Discussion

### A Protocol for Water-in-oil Emulsion Analysis Using Oil-based Flow Cytometry – practical sampling considerations

Oil-based flow cytometry systems have not previously been used for droplet analysis. For this reason, the basis of this work was the establishment of a robust protocol for droplet content analysis using oil-based cytometry on the Guava EasyCyte flow cytometry system 6HT-2L. In particular, two practical problems that had previously precluded a reliable analysis of droplet samples had to be addressed:

i. In the heterogeneous biphasic droplet/oil sample a tendency of droplets to float on fluorous oil was observed, making it difficult to obtain a continuous supply of droplets for analysis
ii. Evaporation from droplets during the course of measurements affected their size and had to be minimized to obtain data faithfully representing the original state of the water-in-oil droplets.

Aqueous droplets have a lower density than fluorinated oil, making them float to form a raft when left undisturbed. Since the sample needle of the flow cytometer is positioned near the bottom of the sample in a 96-well plate, droplets that have floated to the top of the oil phase are not initially drawn in and measured. When the volume in the sample well is nearly depleted, the floating droplets get close enough to the sample needle to be drawn in and measured. The degree to which the floating of droplets impedes sample measurement correlates inversely with oil viscosity: low viscosity oils allow rapid floating, causing the droplets to be poorly sampled at the beginning of the measurement, because they sit at the top of the oil, out of reach of the sample needle. In contrast, when an oil with a higher viscosity, such as FC-40, is used, droplets are detected at the beginning of the measurement (∼ initial 20 seconds), before they have settled, and then they are again detected at the end of the measurement, once the sample volume is depleted (Fig. 2A & Fig. S1, Supporting Information, SI). Such interrupted detection leads to the appearance of two distinct populations in the forward *vs.* side scatter plot (Fig. 2A). The population detected towards the end of the measurement is shifted to lower forward scatter values (FSC). Microscopic examination of the droplets incubated in the well indicated that the droplets had reduced in volume (Fig. S2, SI). We then sought to confirm that the observed second population was indeed caused by droplet shrinkage and to check whether their reduced size led to significant changes in any flow cytometry parameters other than forward scatter.

**Figure 2.**
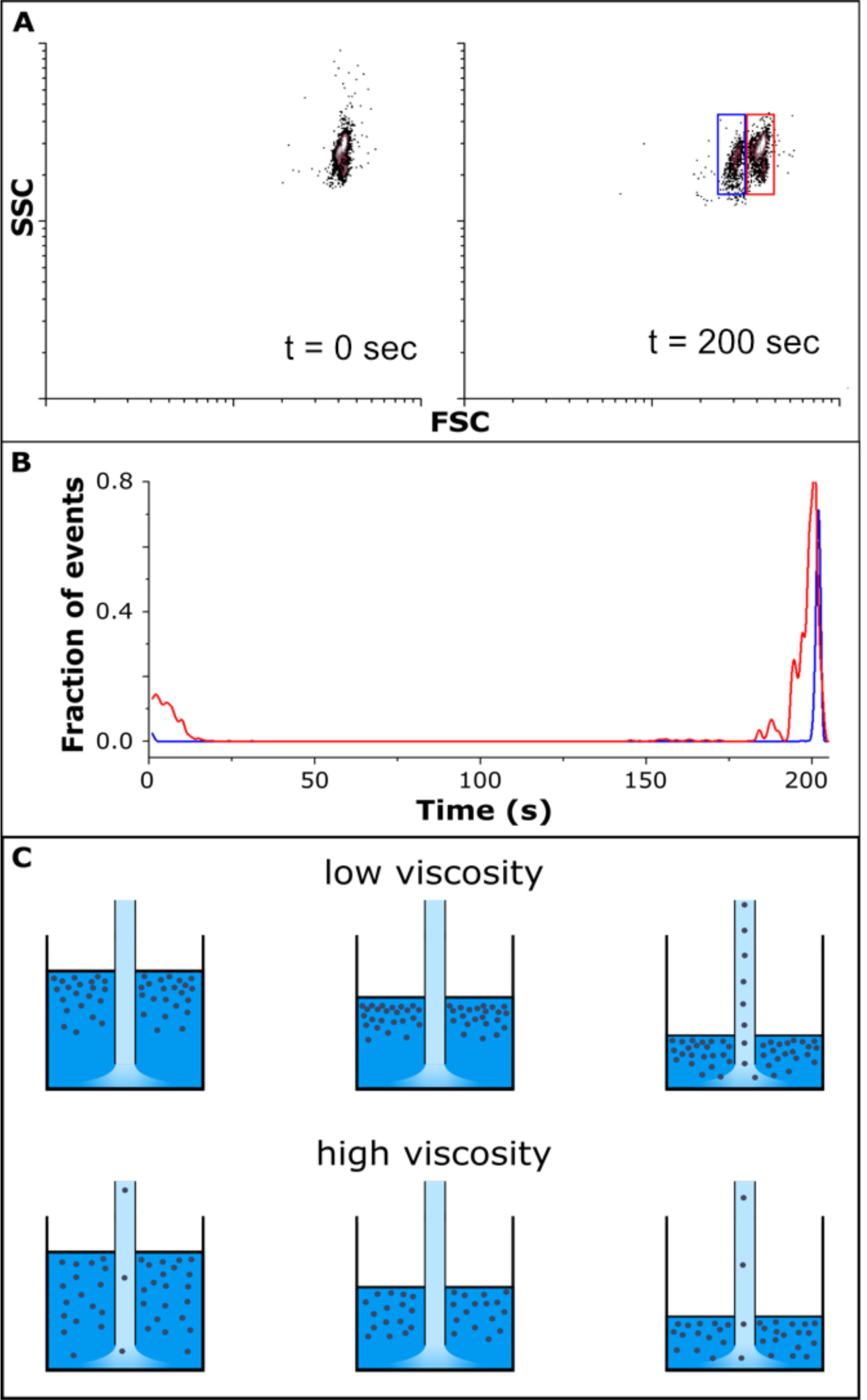
Practicalities of a new small molecule leaking assay. The key experimental consideration was how to supply sample to the flow cytometer. The sample is sucked into the flow cytometer with a nozzle from a microwell during ∼200 seconds. **(A)** Flow cytometric analysis of droplets containing the fluorophore fluorescein **3**, just one population was observed in a scatter plot (logFSC-logSSC; FSC: forward scatter, SSC: side scatter), but subsequently a second droplet population at lower FSC values (*right*, blue box) appears in addition to the first one (red box), consistent with evaporation of the aqueous phase. **(B)** The variation of droplet density of the sample as it is sucked into the nozzle suggests inhomogeneous distribution of droplets in the sample. The initial population (framed in red) shows up when a proportion of droplets dispersed in oil is sampled and again when droplets floating at the surface are sampled. The blue population does not appear continuously, but arises late (generated by evaporation and floating) and is also sampled when the surface nears the nozzle. The x-axis represents the time of the measurement and the y-axis the relative proportion of droplets detected at a given time point in relation to the sum of all droplets. **(C)** The observed distribution of FACS events over the time of sampling depends on the propensity of droplets to float in a given oil (and thus on oil viscosity). *Conditions*: T = 25 °C; droplet content: 100 mM Tris/HCl, 100 mM NaCl, pH 8.0, Oil: HEE-7500 with 1% AZ900C (w/w).

To test the effects of droplet volume reduction on flow cytometric parameters, droplets containing the fluorophore fluorescein **3** were shrunk by osmosis through incubation with fluorophore-free high-salt droplets (0.5 M NaCl).^57–58^ This procedure mimics droplet evaporation in a 96-well plate prior to flow cytometric analysis. When the droplet diameter decreased by more than 2-fold, a 10-fold reduction in forward scatter (FSC) was observed, contrasting with decreases in side scatter (SSC) and green fluorescence (GRN) values of just 1.5-fold and 2-fold, respectively (Fig. S3, SI). These data suggest that evaporation is not likely to have a substantial effect on fluorescence accuracy, at least for measurements not exceeding 15 minutes (Fig. S4, SI). However, the reduced FSC might cause the droplet population to fall below the detection threshold, if the forward scatter threshold had been set too high at the stage of initial gating. Thus, the possibility of droplet shrinkage should be taken into account when selecting the forward scatter threshold, and we recommend that the threshold be set some distance below the droplet population.

Thus, while the Guava EasyCyte instrument is able to automatically sample each well of a 96-well plate, this feature is not helpful for droplet analysis due to droplet evaporation. Instead we developed an alternative procedure, in which samples were transferred into a 96-well plate and mixed immediately before the measurement. With this procedure, the use of sheath fluid-free flow cytometry for droplet analysis allowed very fast and simple measurements consuming very low sample volumes (∼5 µL of emulsion that is typically highly viscous and difficult to handle). It is important to note that oil-based flow cytometry systems are useful for analytical purposes only, as *sorting* of the desired hits is impossible using this instrumental set-up. Droplets can be sorted on-chip^16–17, 26–27^ or, after forming double emulsions^52–56^ or gel-shell beads,^59^ in a FACS instrument, but the information gained from flow cytometric analysis as described here provides the basis for setting up either of these sorting experiments, and offers a ‘quality control’ step. The ability to quickly assess the feasibility of a droplet assay and set up optimal conditions is extremely valuable to avoid the loss of precious library sample and will make droplet-based experiments more accessible, by rapidly providing feasibility information.

### Quantification of Small Molecule Leaking

Minimization of small molecule transfer is crucial to expanding the variety of reactions that can be assayed in microdroplets and the accessible timescales over which reaction progress can be faithfully monitored. We chose resorufin **1** as a representative small fluorescent molecule, because it is the product of many commercially available enzyme assays (e.g. a leaving group in hydrolytic reactions^26, 60–61^ or a redox indicator in the Amplex reagent family^25, 62–65^). Resorufin **1** transfers rapidly between droplets and we set out to quantify small molecule transfer by mixing fluorescent resorufin-containing droplets with non-fluorescent, buffer-containing droplets, using a sheath fluid-free flow cytometry for droplet analysis. Oil-based flow cytometry systems have not previously been used for droplet analysis and we provide the first study, using a Guava EasyCyte instrument, of the convergence of two, initially distinct, populations over time.

To facilitate the interpretation of our results we introduce a parameter that quantifies the amount of fluorescence that has transferred between droplets at any particular time. This parameter (’% leakage’) is defined by equation 1:

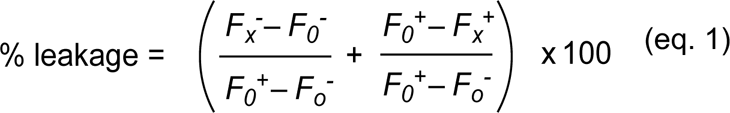

where 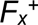 is the mean fluorescence value of droplets originally containing the fluorophore (‘positives’, superscript “+”), and 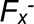 refers to those that did not (‘negatives’ superscript ‘-’) in populations at times 0 or x (denoted by subscripts 0 or x), as determined by flow cytometry. This equation thus describes the degree to which a sample has progressed from the initial clear separation towards homogeneity (Fig. 3).

**Figure 3.**
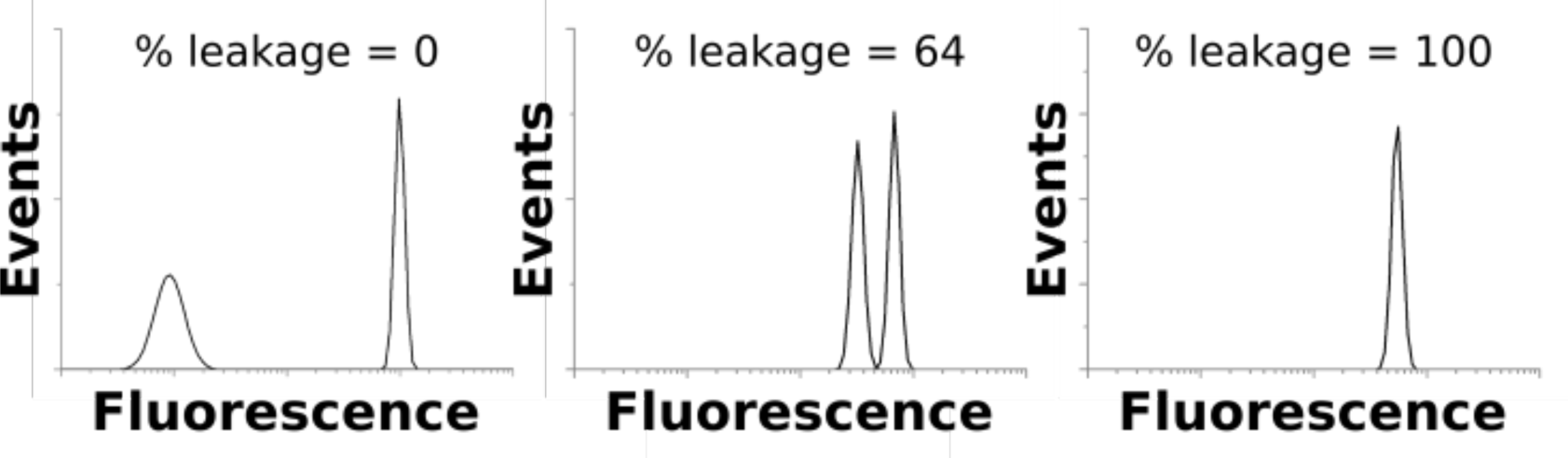
Distribution functions of events (i.e. droplets passing the detector) monitored in a flow cytometric gate and their corresponding representative % leakage values. Initially strongly fluorescing (superscript “+”) and non-fluorescent populations are clearly separated (left panel). Leakage then renders the fluorescence values increasingly similar (corresponding to peaks that are separated less clearly, middle panel), until they finally merge into one peak representing homogeneous distribution of the fluorophore in all droplets (right panel). For this figure example data (obtained with resorufin **1,** FC77 and 1% AZ900C) were used to illustrate the correlation of fluorescence distribution functions and the corresponding % leakage values (calculated using equation 1).

### Quantitative comparison of retention of different fluorophores

To exemplify the utility of our analytical approach leakage rates were measured for four commonly used fluorophores (Scheme 1). Figure 4 shows the time courses starting from initially separate populations containing the fluorophore or not. In three cases the first measured time point shows these populations as still distinctly separate in flow cytometric analysis, while for rhodamine B (**4)** leakage is so fast that convergence to one peak has already occurred in the ‘dead time’ of the protocol (< 1 min). The time courses resulting from calculation of % leakage was calculated (eq. 1) from the primary data in Fig. 4A show that retention half lives range from <1 min (**4**), to ∼37 min and ∼94 h (for **1** and **3**, respectively), with fluorescein **3** being almost completely retained.

**Fig. 4.**
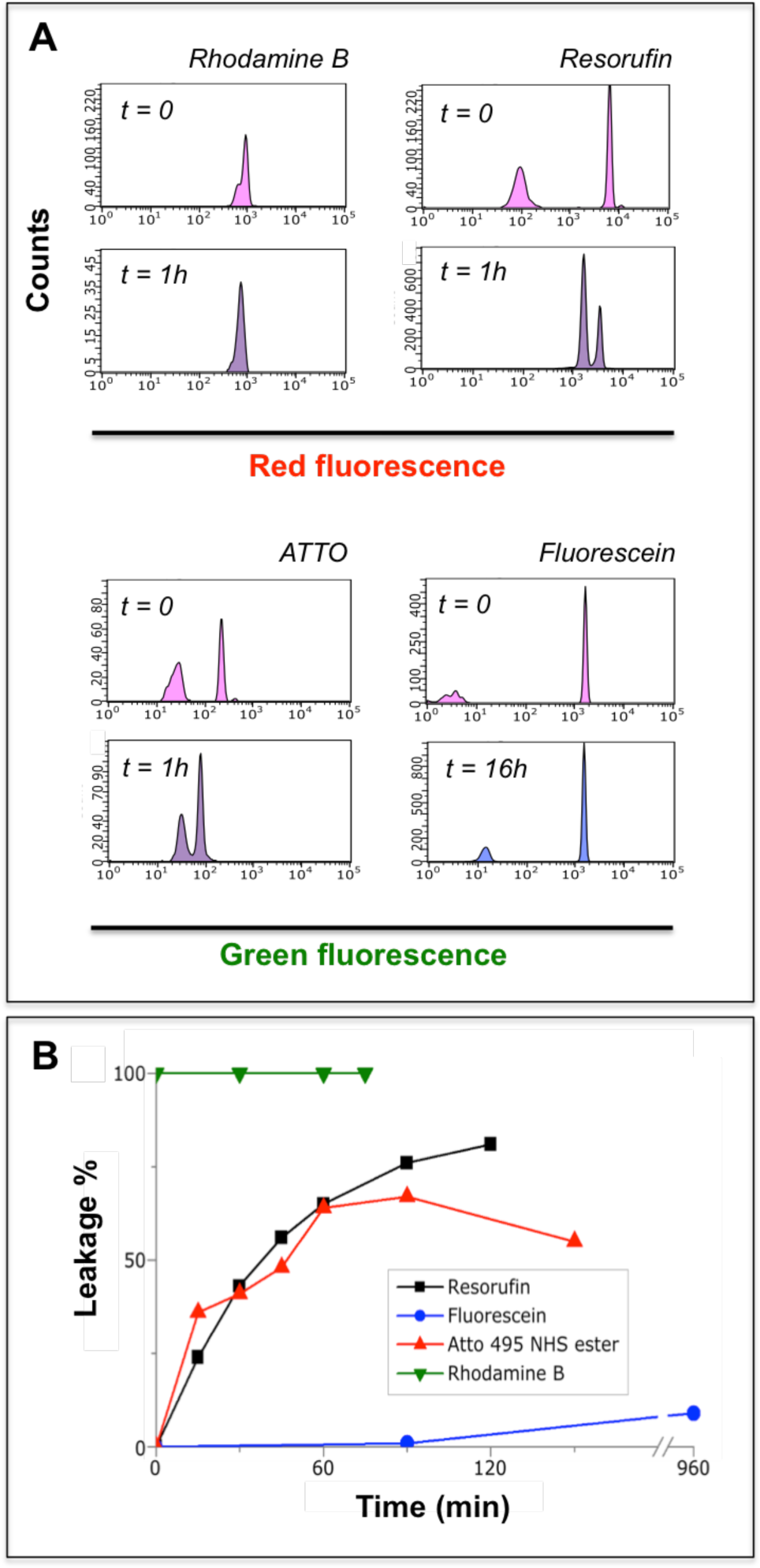
Time courses of small molecule transfer for different fluorophores. To exemplify the utility of our analytical system leakage rates were measured. **(A)** Primary flow cytometric data of measured to determine a time course. Two samples – of non-fluorescent and, on the other hand, highly fluorescent droplets - were mixed in an Eppendorf tube (at time point zero, t=0) and measured shortly afterwards (10-20 s). Initially separate peaks eventually converge (in the case of rhodamine B already within the time taken to obtain the first measurement). **(B)** The % leakage values were calculated using eq. 1 from the primary data in (A) and are shown as a function of time for the four fluorophores examined. *Conditions*: 25 °C, HFE7500 with 1% AZ900C (w/w).

**Scheme 1.**
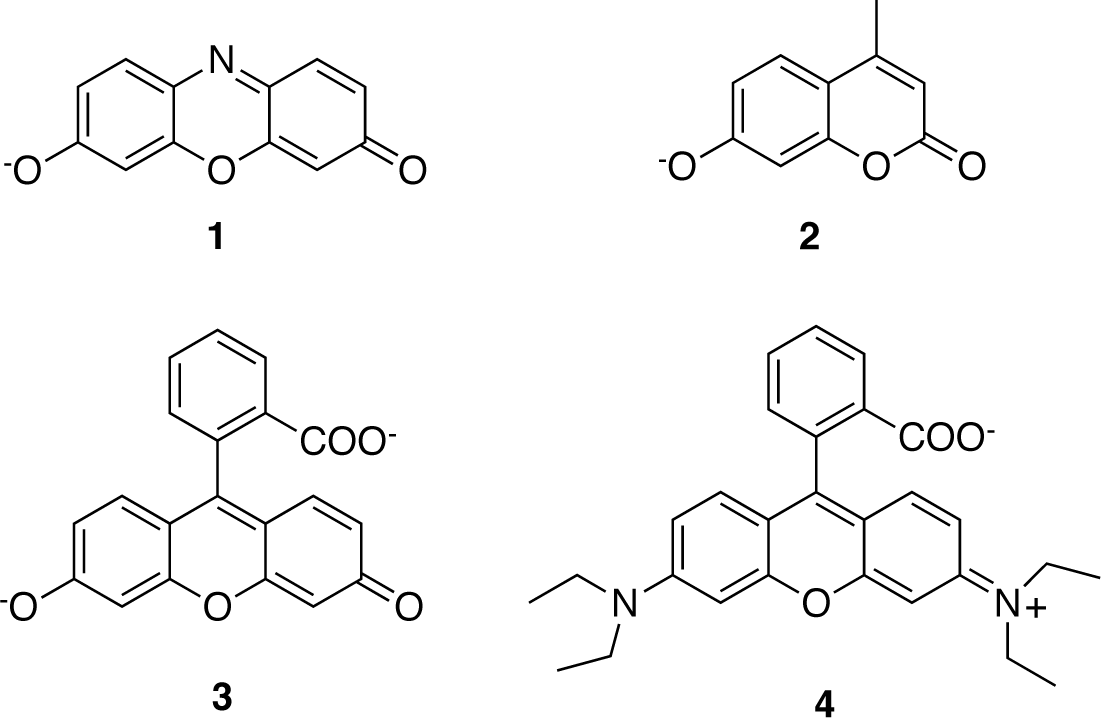
The fluorophores used in this study were resorufin **1**, 4-methylumbelliferone **2**, fluorescein **3** and rhodamine B **4.** The molecules are shown in their deprotonated state that is present at pH 8 and represent the fluorescent species.

### Ionisation of the Fluorophore Increases Retention

Prompted by a previous report in which the poorly-retained fluorophore coumarin was chemically derivatised (with a methylsulfonate group, introducing a negative charge) resulting in more than a 90,000-fold reduction in leakage,^35, 66^ we investigated the pH dependence of leakage for the ionizable phenolate resorufin **1** (Fig. 5; Table S3, SI) and found that the pH of the aqueous solution within droplets has a major influence on fluorophore retention. Higher pH improved retention of resorufin **1**, presumably due to the greater extent of deprotonation, reducing the driving force for partitioning into the oil phase and slowing diffusion. It appears that the p*K_a_* of resorufin (in solution: 5.8)^61^ is slightly perturbed in the droplet (or at the water-oil interface) as the pH has to be raised well above the p*K_a_* to show the improvement. At pH 9.0 resorufin **1** leakage was slowed more than 40-fold, allowing two distinct fluorescent populations to be identifiable for up to two hours, while at pH 6.0 leakage was so rapid that two initially distinct fluorescent populations converged during the 3 minute measurement of a single flow cytometry run (Fig. S6, SI).

**Figure 5.**
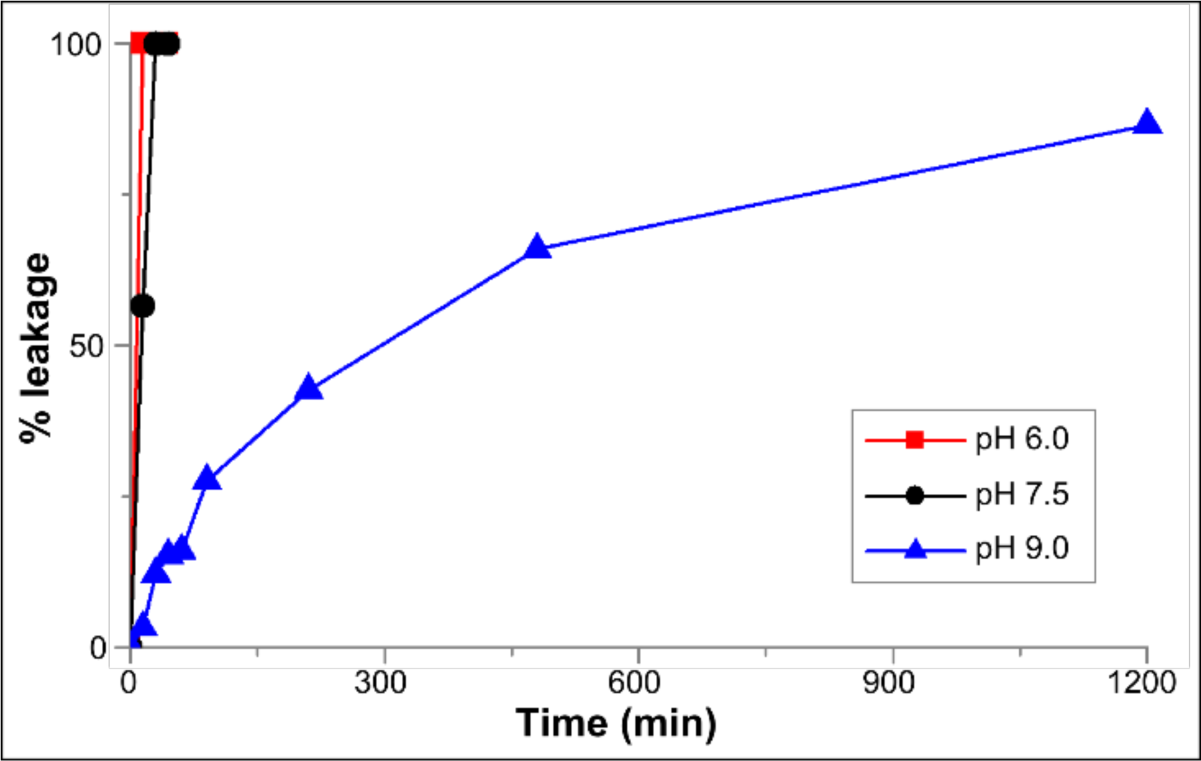
Factors effecting leakage of resorufin **1** from droplets. Time courses of % leakage were obtained from flow cytometric measurement (as shown in Figs. 3 & 4) Droplets with an aqueous phase with higher pH retain ionisable fluorophore molecules up to 1200-fold longer than those with a lower pH values, suggesting a correlation of small molecule charge and leakage: the increasing deprotonation of resorufin **1** at higher pH leads to better retention. *Conditions*: T = 25 °C, droplet content: 100 mM Bis-Tris propane, 100 mM NaCl, pH 6.0, 7.5 or 9.0, respectively, with addition of 10 µM resorufin **1** for positive droplets. Oil: HEE-7500 with 1% AZ900C (w/w).

### The Role of the Fluorinated Carrier Phase: Oxygen-containing Oils Facilitate Small Molecule Transfer

Transport into and through the oil phase is one of the main mechanisms of small molecule transfer between water-in-oil emulsion droplets. We examined the rate of fluorophore transfer between droplets for a selection of commercially available fluorous oils (covering different chemical structures and physical properties). The oils we used can be divided into four structural groups (Fig. 6A). HFE oils consist of perfluoroalkyl chains of varying lengths that bear an alkoxy group (i.e. methoxy or ethoxy). The ‘FC oils’ can be subdivided into three groups. First are the simple perfluorinated linear alkyl oils FC72 and FC77, the second group of FC oils are the amine-centered oils FC3283, FC40, FC43, and FC70 which each contain a tertiary amine bearing perfluorinated alkyl chains of different lengths. FC40 is a physical mixture of FC43 and di(perfluorobutyl)methylamine. Finally, FC3284 is a morpholine-derived oil that contains both a tertiary amine and an oxygen atom in an ether functionality. A summary of the physical properties of these oils is provided in the SI (Table S1).

**Figure 6.**
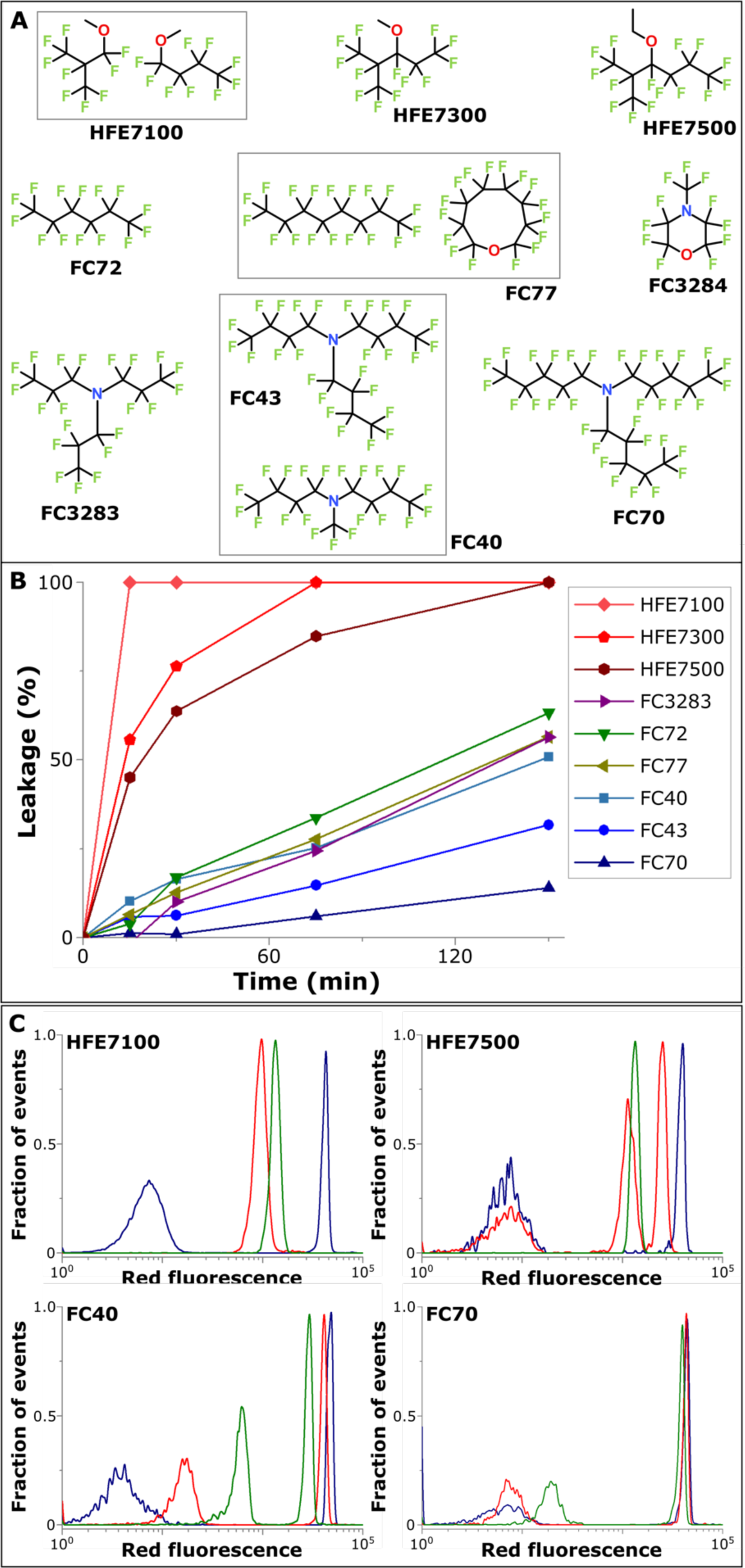
The effect of the carrier oil on the rate of leakage of resorufin **1**. **(A)** Molecular structures of fluorinated oils tested. **(B)** Time courses of leakage over 2.5 hours for a series of the oils shown in (A). HFE-type oils are worst at small molecule retention. The retention of small molecule by FC-type oils improves with increasing viscosity (see Table S1, SI). **(C)** Superposition of FACS histograms depicting fluorescence for chosen oils from each structure class as a function of time. For each graph the time points presented are 0 min (blue), 30 min (red) and 150 min (green). HFE-7500 and FC-40, the most commonly used fluorinated oils in the field microfluidics, are compared with another type of oil from the same class with a lower (HFE-7100) or higher viscosity (FC-70), respectively. *Conditions*: T = 25 °C, Droplet content: 100 mM Tris/HCl, 100 mM NaCl, pH 8.0 with addition of 10 *μ*M resorufin **1** for positive droplets. Oil: HEE-7500 with 1% AZ900C (w/w) for droplet formation, exchanged into oils of interest containing 1% AZ900C (w/w).

Monodisperse droplets with and without resorufin **1** (10 μM) in buffer were produced in a flow focusing chip (see Fig. S8, SI), with HFE-7500 as the carrier oil. HFE-7500 was used for droplet production as its low viscosity allows rapid production (> 10 kHz) of monodisperse droplets. Aliquots (50 µL) were then transferred into the oils tested (500 µL) and small molecule transfer was monitored by flow cytometry over time (Figs. 6B,C). A wide range of small molecule transfer (% leakage ranging between 12 and 100% after 150 minutes) was observed. The small molecule transfer is most rapid for the HFE-type oils, such that after 15-60 minutes of incubation small molecule transfer is already almost complete (Figs. 6B,C; Table S2, SI). With the exception of FC3284, the FC-type oils show significantly greater retention of resorufin **1** than HFE oils (data not shown). Among the FC-type oils, the oils FC-72 and FC-77 (that are predominantly fluorocarbon oils) show lower retention of resorufin **1** than the amine-containing oils. FC3284 leaks rapidly, similarly to the oxygen-containing HFE oils. The chemical structure of the oil thus appears to be the primary determinant of its ability to retain resorufin **1**: empirically, oils that contain an oxygen atom leak rapidly, those with no heteroatoms leak comparatively slowly, and oils containing a nitrogen atom leak most slowly. However, there is no obvious molecular explanation for this effect. Plotting leakage against a variety of physical properties of the tested oils did not reveal direct clues to properties that could be optimized to reduce small molecule transfer further (Figure S5, SI). It was noted, however, that within each structural class small molecule transfer tended to decrease as viscosity, molecular weight and boiling point increased. These physical properties are interdependent: higher molecular weight compounds are more viscous and have higher boiling points.

Finally, droplet stability (i.e. maintaining the clonal compartmentalisation for an experimental workflow that involves the process of compartmentalisation to optical interrogation of droplets and sorting) is an important factor when selecting which oil to use. We have observed that some FC-type oils, despite providing a 4- to 60-fold greater retention of resorufin **1** than HFE-type oils, stabilize droplets less well than HFE-type oils (i.e. coalesce within hours in experiments that require droplet handling and incubation).^29^ The difference in stability becomes particularly evident when working with challenging samples such as cell lysate assays in droplets. Cell components, as well as detergent-containing cell-lysis agents, interact with droplet-stabilizing surfactant molecules and counteract droplet stability. Here, we implement a new approach to managing these divergent effects by forming droplets in an HFE-type oil that provides good stability and subsequently exchange the sample into a leakage-minimizing FC-type oil.

### Small Molecule Transfer is Affected by the Nature and Concentration of the Surfactant

The identity of the surfactant is known to affect the stability of droplet formulations, but, while retention data are available for mineral oils/hydrocarbon surfactant pairs, leaking studies are available for fluorous systems. We determined time course of leakage of resorufin **1** from droplets and compared this process in droplets stabilised by the commercially available surfactants Pico-Surf 1 (structure and composition not known) and 008-FluoroSurfactant (likely to be similar to the surfactant described by Holtze *et al*.^45^) and the new surfactant AZ900C (see section D, SI, for the synthesis). A clear ∼4-fold difference in retention half-lives is evident, with Pico-Surf 1 loosing the fluorophore more quickly than the other two (Fig. 7).

**Figure 7.**
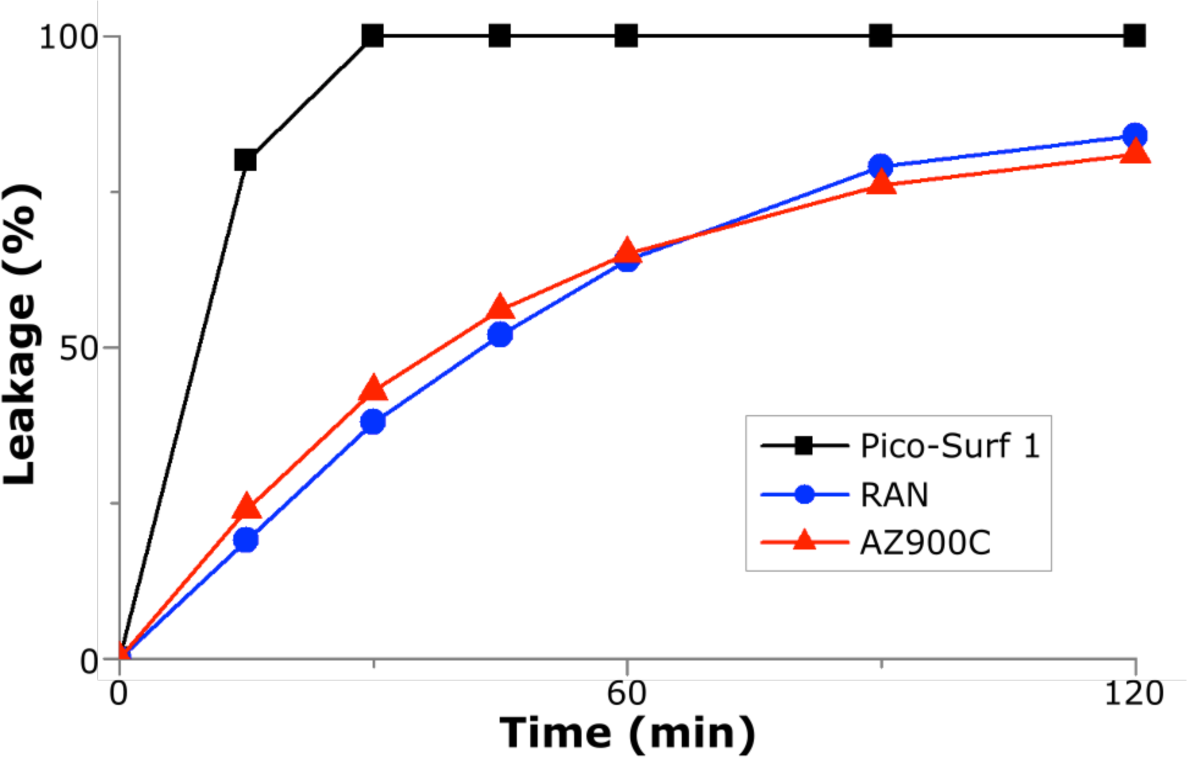
Leakage depends on the surfactant type. The fluorophore resorufin leaks ∼4-fold faster from droplets stabilised by the commercially available surfactant Pico-Surf 1 (Dolomite) than from those maintained by 008-FluoroSurfactant (Ran Biotechnologies) or the new surfactant AZ900C (see section D, SI). *Conditions*: = 100 mM Tris/HCl, 100 mM NaCl, pH 8.0 with addition of 10 µM resorufin **1** for positive droplets. Oil: HEE-7500 with 1% (w/w) Pico-Surf 1, 008-FluoroSurfactant or AZ900C.

Next we tested the effect of different concentrations of surfactant using the two most commonly used fluorinated oils – HFE-7500 and FC-40, each representing one of the structural classes (see below). An inverse relationship between the retention of resorufin **1** in droplets and surfactant concentration was observed (Fig. 8A, Table S2, SI). HFE-7500 facilitates small molecule transfer to a much higher extent (6- and 14-fold more for 1 or 0.1% surfactant, respectively) than FC-40. This means that small molecule transfer can be attenuated by variation of surfactant concentration: for HFE-7500 a decrease from 1% to 0.1% leads to better separation between the positive and negative populations for up to 60 minutes (Fig. 8B). It has been suggested that surfactant molecules mediate the solubility of fluorophores in the fluorinated phase.^67^ The best retention of resorufin **1** in droplets was achieved using 0.1% surfactant AZ900C in FC-40 (30% less leakage than FC-40 with 1% AZ900C after 3 h). Based on the observed improvement in fluorophore retention at low surfactant concentration, it would be tempting to reduce the surfactant concentration even further, but this is not possible as the surfactant is responsible for droplet stabilization. Decreasing surfactant concentration too much can greatly reduce droplet stability and lead to droplet merging, which we observed particularly for samples containing surface-active ingredients, such as dense cell lysates or lysis agents. While merged droplets can be easily ‘gated out’ for purposes of analysis, droplet merging would nullify the compartmentalisation of individual experiments, in which each droplet contains e.g. a different set of reagents (e.g. members of a library of DNA, protein or small molecules or simply various buffers or additives) and jeopardise selection. In our experimental design it proved crucial to strike a balance between droplet stability and minimization of small molecule transfer through careful adjustment of surfactant concentration for the individual sample. Oil exchanges between steps with different requirements provide means to take advantage of the properties of multiple oil formulations sequentially.

**Figure 8.**
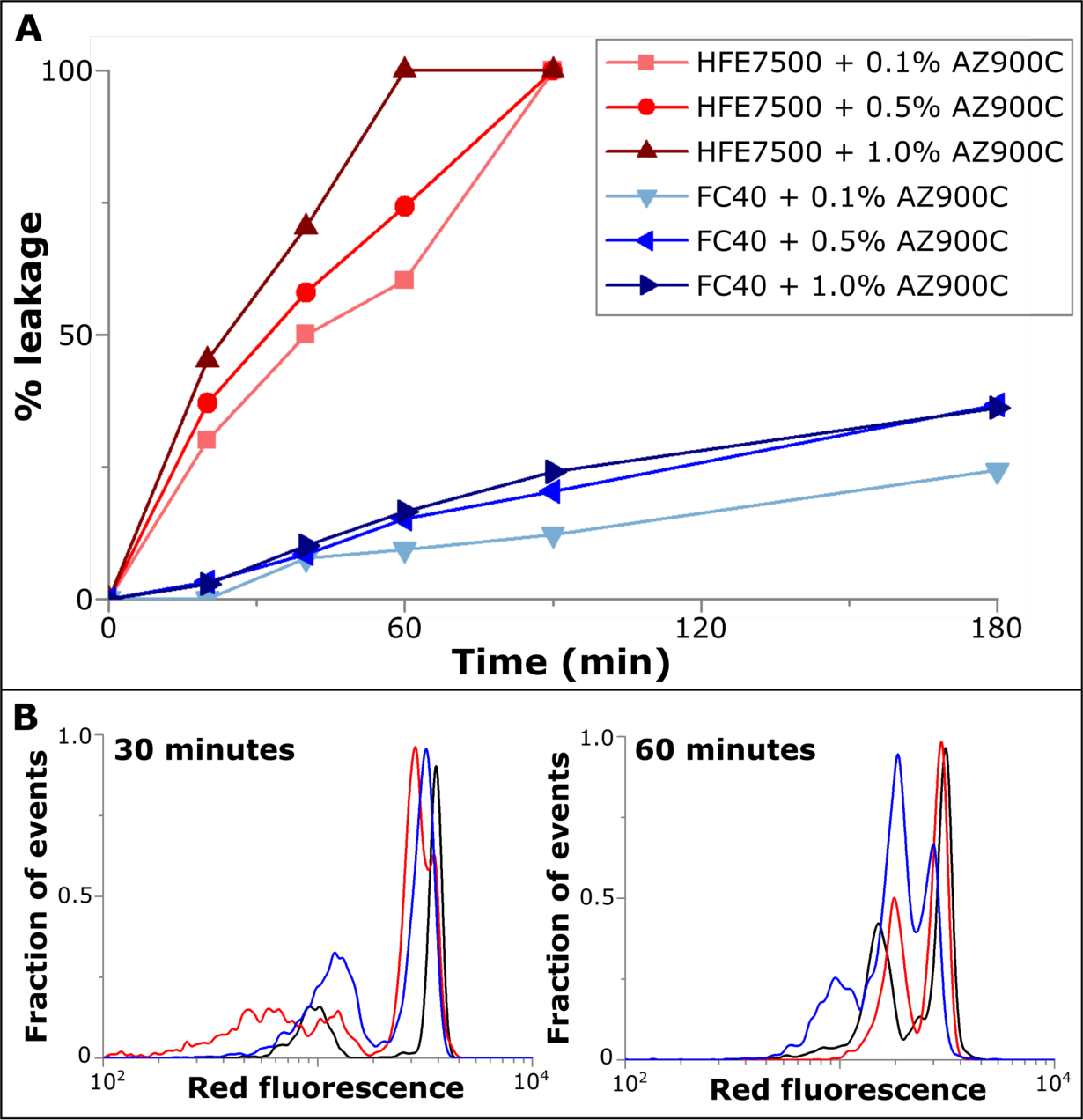
Leakage depends on surfactant concentration as well as oil type. **(A)** % leakage (calculated according to eq. 1) over a time period of 2.5 hours for the range of oils examined. HFE-7500 shows much greater leakage than FC40, but the leakage of both oils is reduced when the concentration of surfactant AZ900C is lowered. **(B)** Overlay of fluorescent populations after varying incubation times (30 minutes, *left*; 60 minutes, *right*) in HFE-7500 with 0.1% (red), 0.5% (black) and 1% (blue) surfactant (w/w). These histograms show that when low surfactant concentration is used the leakage of even HFE-7500 is reduced sufficiently to allow distinction of positive and negative populations after as long as 60 minutes. *Conditions*: T = 25 °C, droplet content: 100 mM Tris/HCl, 100 mM NaCl, pH 8.0 with addition of 10 *μ*M resorufin **1** for positive droplets. Oil: HEE-7500 with 1% AZ900C for droplet formation, exchanged into oils of interest containing AZ900C concentrations as indicated in the box.

### Aqueous Phase Additives Improve Retention

Addition of BSA to the aqueous phase has been reported to reduce small molecule transfer by blocking the interface between the oil and the aqueous phase or by changing the partition coefficient by small molecule binding.^33–34^ The combination of optimized conditions derived above (FC-70 with 0.1% surfactant AZ900C) with 1% BSA added to the aqueous phase enabled detection of two well-resolved fluorescent populations for up to 130 hours (Fig. 9). This is a more than 40-fold improvement compared to previously reported conditions (HFE-7500 with 0.5% KryJeffa surfactant), which resulted in complete leakage after 3 hours.^34^ This substantial reduction in resorufin **1** leakage dramatically increases the range of assay timescales, so that even assays using slow enzymes that require long incubation times to produce sufficient optical signal become feasible. The ability to prolong retention raises the possibility of using the many commercially available substrates derived from resorufin **1** for enzymatic assays in microfluidic droplets that, due to their rapid leakage, were hitherto considered useless for droplet-based experiments.

**Figure 9.**
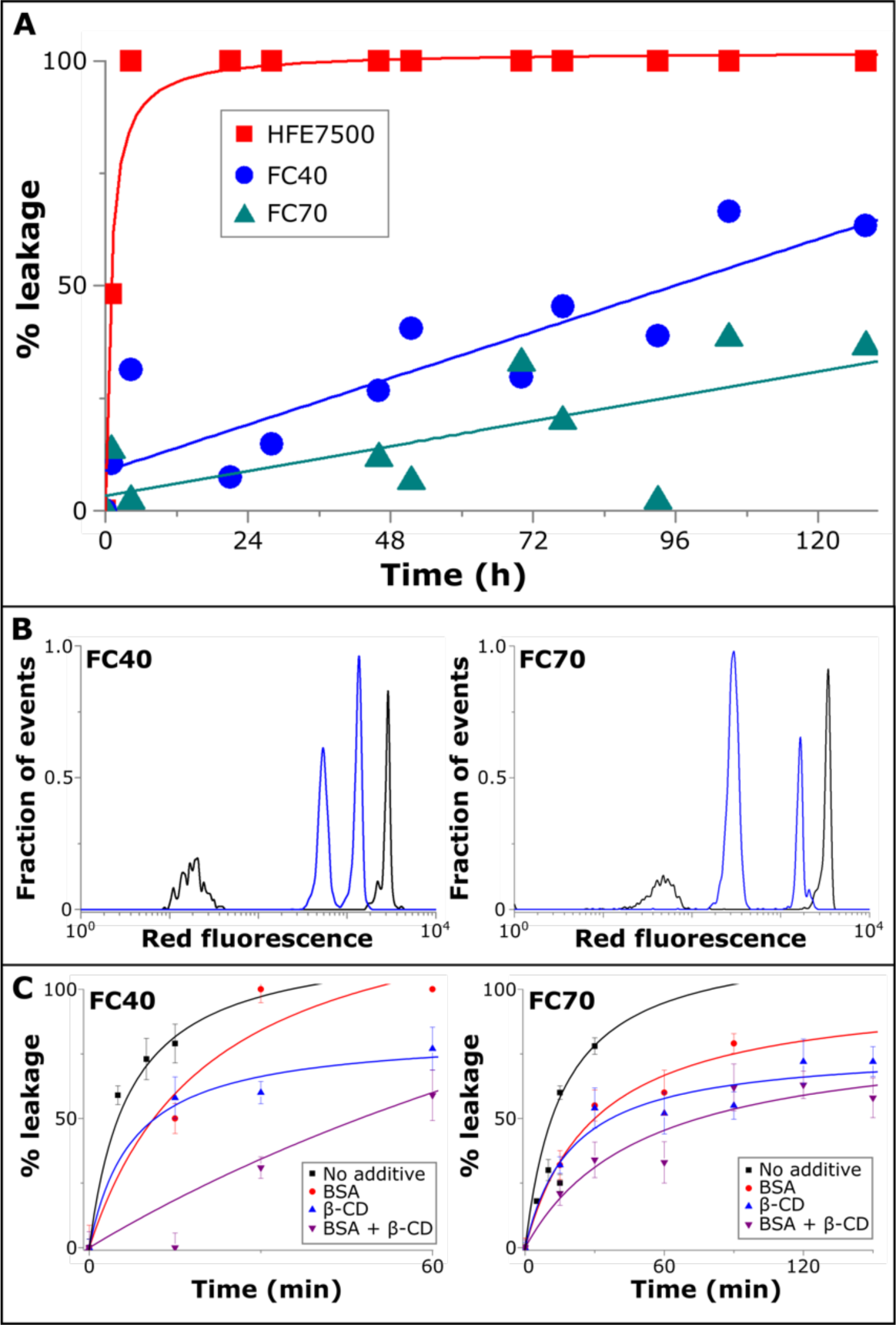
The aqueous-phase additives BSA and β-cyclodextrin enhance the retention of fluorophores. **(A)** Influence of the addition of BSA (1%, w/w) on the leakage of resorufin **1** over a period of 5 days. The curves were merely drawn to guide the eye. **(B)** Peak separation of initially-distinct fluorescent populations after 26 hours or 43 hours in FC40 or FC70, respectively, in the presence of BSA (1%, w/w) in the aqueous phase. **(C)** Retention of 4-methylumbelliferone **2** in the presence of aqueous phase additives BSA (1%, w/w) or β-cyclodextrin (10 mM) or both additives) in FC40 or FC70, respectively, as determined by analysis of microscopy images. *Conditions*: T = 25 °C, Droplet content: 100 mM Tris/HCl, pH 8.0, 100 mM NaCl, with addition of 10 *μ*M resorufin **1** or 20 µM 4-methylumbelliferone **2** for positive droplets. Oil: as indicated, with 0.1% AZ900C.

Despite this achievement, the retention of other small hydrophobic fluorescent molecules, such as 4-methylumbelliferone **2** (the product of widely used fluorogenic assays for alkaline phosphatase, β-galactosidase and β-glucuronidase among others),^68–72^ continues to be challenging. We investigated whether BSA could aid in the retention of 4-methylumbelliferone **2** in droplets and found that it was able to reduce leakage 3-fold (a 3-fold smaller effect than that seen for resorufin **1**, Tables S2 and S4, SI).

In addition, we then tested whether complexing agents could help in retention of small molecules. β-cyclodextrin is a well-known complexing agent for hydrophobic compounds,^73^ so its effect on retention of 4-methylumbelliferone **2** was tested. Using both FC-40 and FC-70 with 0.1% surfactant AZ900C and an excess of β-cyclodextrin (10 mM *vs* 20 µM 4-methylumbelliferone **2**) as an additive, we succeeded in extending the retention time 15-fold to 150 minutes compared to a control experiment without β-cyclodextrin (Fig. 9C). The leakage in HFE7500 was very rapid (complete in less than 6 min; Fig. S7, SI) even when both leakage-slowing additives (1% BSA and 10 mM β-cyclodextrin) were present in droplets. Without additives the leakage in HFE-7500 is barely measurable (complete within 1-2 minutes), 75-fold faster than under optimized conditions (see above) that enable an extension of the fluorophore retention time to 150 minutes. The effects of BSA and β-cyclodextrin were roughly additive: a combination of both improved the retention of 4-methylumbelliferone **2** beyond the effect of β-cyclodextrin alone (Table S4, SI). Despite the greatly improved retention of 4-methylumbelliferone **2**, leakage remains relatively rapid, meaning that 4-methylumbelliferone **2**-based substrates should only be used for fast reactions that can be incubated and sorted on chip within two hours. However, by increasing its retention time to at least 2.5 hours (see Fig. 9C), our optimized conditions (10 mM β-cyclodextrin, FC-70 with 0.1% surfactant AZ900C) provide a sufficient time window for double emulsion formation on a second microfluidic chip followed by FACS sorting.^52^

## Conclusions

The availability of this new, simple, user-friendly method for flow cytometric analysis of water-in-oil emulsions will provide a reliable quantification of the time window in which droplet measurements involving small molecules can be carried out without being compromised by leakage. The utility of this approach is demonstrated by an enlargement of this window for leakage of the fluorophore resorufin **1** by more than an order of magnitude as a consequence of careful optimization of the oil, surfactant and aqueous phase properties guided by the results obtained using our flow cytometric analysis.

General lessons can be learned by systematic analysis of the influence of oil type, surfactant concentration, pH within droplets and utilization of additives for retention of small fluorophore molecules in droplets. Consistently an FC-type oil is best (with FC-70), and a low surfactant concentration (0.1%), and a pH at which the fluorophore is ionised lead to the best retention. The emerging best set of conditions to retain resorufin **1** in droplets (FC-70 with 0.1% w/w surfactant AZ900C) makes clear improvements, resulting in at least 20-fold longer retention time than previously described (based on the comparison of the times taken for 50% leakage, i.e. 0.3 h (or 0.5 h in ^34^) *vs* ∼10 h extrapolated from the data in Fig. 6). Addition of 1% BSA to the droplet led to a further increase of retention time, enabling resorufin **1** retention for at least five days, representing an improvement in retention of over 40-fold. These guidelines can be generalized to other small molecules: the retention of 4-methylumbelliferone **2**, a fluorophore that is extremely prone to leakage into neighboring droplets was improved by at least 30-fold (under the same conditions established for optimal retention of resorufin **1** in combination with the complexing agent β-cyclodextrin). Because of the low solubility of organic molecules in fluorinated solvents, molecule concentration in the fluorinated phase can be neglected.^67^ Practically, we devise a procedure in which the properties of different oils can be sequentially exploited: HFE-type oils provide good stability and are the best choice for the droplet formation step, while – after an oil exchange step - leakage-minimizing FC-type oils should be used for long-term incubation.

Previous success in improving the retention in microdroplets of leaky fluorophores was achieved by chemical modification of the fluorophore itself. While chemical modification of the fluorophore leads to more dramatic changes in retention than optimization of oil, surfactant and aqueous additives, it can in many cases be synthetically non-trivial, prohibitively laborious and/or costly, and has the significant disadvantage of altering the molecular assay being studied. Furthermore, any chemical modification will need to be optimized independently for each fluorophore.

The reductions in leakage achieved here will usefully broaden the scope of droplet assays to include slow reactions, thus rendering the microdroplet format suitable for reaction that may require an extended time window: sorting of metagenomic libraries^22, 74^ and libraries for directed evolution of typically slow promiscuous enzyme activities^75–76^ or for enzymatic assays of low abundance receptors with single cells in droplets,^13^ but also for small molecule enzyme inhibition studies.^77–79^ Our analytical approach is simple to apply, enabling ready monitoring of fluorescence of droplets and will complement mass-spectrometric analysis ^47^. Rapid assessment of leaking will be useful to researchers looking to monitor the progress of microdroplet trial-reactions in established systems as well as in cases where new reactions and conditions for microdroplet experiments are developed. Ultimately a growing empirical body of evidence will contribute to the development of guidelines that are broadly applicable to improving the retention of any small molecule of interest.

## Materials and Methods

### Droplet Generation

Monodisperse water-in-oil-emulsion droplets were formed as previously described.^17, 52, 59, 80–81^ In short, hydrophobic flow-focusing devices with channel dimensions of 20 µm height/20 µm width were operated with flow rates of 100 µL/h total for the aqueous phase and 1000 µL/h for the carrier oil phase, resulting in monodisperse droplets with an average diameter of 18 µm (∼3 pL volume).^52^ The carrier oil phase was HFE 7500 (3M) with 1% (w/w) surfactant (AZ900C, synthesized as described in the SI).

### Flow Cytometry

Flow cytometry measurements were performed with a Guava EasyCyte 6HT-2L using a 96-well microplate following the manufacturer’s instructions. In short, a work list was created by selection of sample containing wells. The mixing function was switched off, as in our experience stirring at the beginning of the measurement reduced data quality, partly by overloading the detector. The total number of events to be acquired was set to 20,000. At the beginning of the measurement, 130 µL of fluorinated oil of choice, such as HFE7500 with 1% surfactant, was added into each selected well, and immediately before measurement 5-10 µL of dense emulsion was added and mixed by pipetting up and down. Before measuring test samples, one run was carried out to adjust settings, with typical values being medium flow rate (0.59 µL/sec), forward and side scatter gains of 1, a threshold trigger set on forward scatter at a value of 300 for experiments with resorufin **1** and on green fluorescence for salt shrinkage experiments (with the green fluorescence gain set on minimum). Red fluorescence (RED-2) gain was adjusted to ensure that the fluorescence of the resorufin **1**-containing samples were on scale. After adjusting settings, the data from this first run was discarded and samples were then run. The automatic sampling function of the cytometer was not used for droplet analysis, but rather the instrument was paused after each sample, the tray was ejected and the next sample was added into the appropriate well of the 96-well plate. Data analysis was performed using GuavaSoft 2.7.

### Assessment of Small Molecule Transfer

To examine small molecule transfer between droplets, two water-in-oil emulsions, one containing fluorophore (10 µM resorufin **1** or 20 µM 4-methylumbelliferone **2** in 100 mM Tris/HCl, 100 mM NaCl, pH 8.0) and another containing only buffer were produced separately in HFE7500 with 1% surfactant as described above. When the effect of additives was examined, both fluorescent and non-fluorescent droplets contained additionally either 1% (w/w) BSA or 10 mM β-cyclodextrin or both additives. Before the first time point measurement equal volumes of fluorescent and non-fluorescent droplets were pipetted into an Eppendorf tube containing a 10-fold volume excess of fluorinated oil–surfactant-mixture of interest. All measurements were taken well above the likely CMC of the surfactants.^32^

### Assessment of Shrinkage

For salt shrinkage measurements, fluorescent droplets contained 10 µM fluorescein **3** in double-distilled water and non-fluorescent droplets 500 mM sodium chloride solution only. Before the first time point measurement, equal volumes of individual droplet species were pipetted into an Eppendorf microtube containing HFE7500 with 1% surfactant (w/w) and mixed.

### Microscopic Imaging of Droplets

In order to ensure that the droplets analysed by flow cytomery retained their integrity before analysis and to quantify retention of 4-methylumbelliferone **2**, images were taken with an Olympus BX51 upright microscope. Fluorescence analysis was performed in the program ImageJ by drawing a line across fluorescent droplets and plotting their grey value profile. The gray value list for each profile was transferred into Microsoft Excel and the average gray value was calculated. Microscope images with >10 droplets were analyzed to quantify 4-methylumbelliferone retention in Figure 9C. Independently it was shown that image analysis of ∼10 droplets yielded equivalent results to analysis of > 50 droplets. The diagnostic parameter ‘% leakage’ was calculated using eq. 1.

## Supporting Information

Supporting Information is available online and contains additional experimental data, a compilation of physical properties of oils and procedures for surfactant synthesis.

## Acknowledgements

This work was supported by the Engineering and Physical Sciences Research Council (EPSRC) and the EU project Metafluidics (685474). FH is an ERC Advanced Investigator (695669). AZ was supported by a BBSRC studentship, the Cambridge European Trust and the EU Marie-Curie network *PhosChemRec*. SRAD received an individual postdoctoral EU Marie-Curie fellowship.

## Table of Contents Graphic

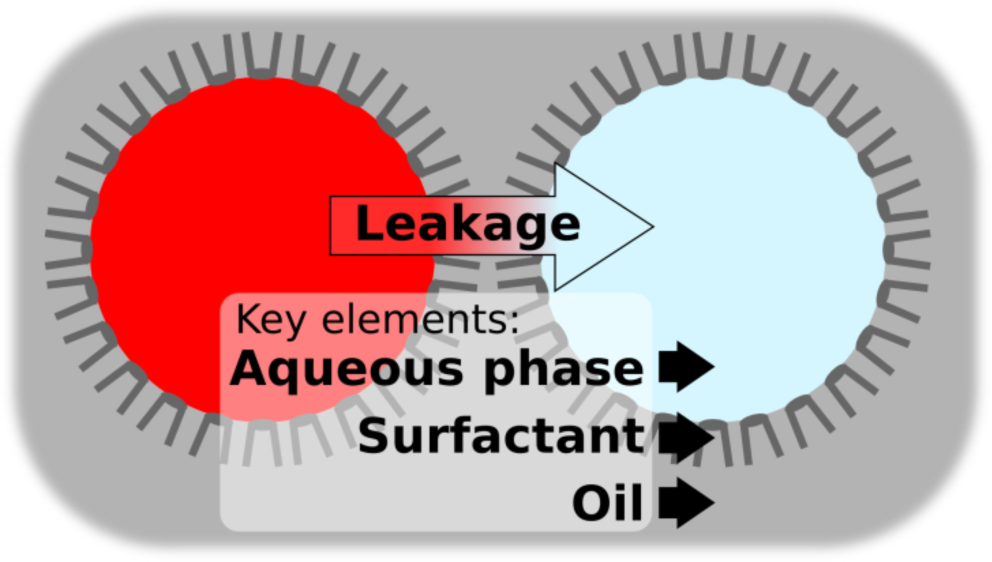

